# Solution structure, glycan specificity and of phenol oxidase inhibitory activity of *Anopheles* C-type lectins CTL4 and CTLMA2

**DOI:** 10.1101/565705

**Authors:** Ritika Bishnoi, Gregory L. Sousa, Alicia Contet, Christopher J. Day, Chun-Feng David Hou, Lauren A. Profitt, Deepak Singla, Michael P. Jennings, Ann M. Valentine, Michael Povelones, Richard H. G. Baxter

## Abstract

Malaria, the world’s most devastating parasitic disease, is transmitted between humans by mosquitoes of the *Anopheles* genus. *An. gambiae* is the principal malaria vector in Sub-Saharan Africa. The C-type lectins CTL4 and CTLMA2 cooperatively influence *Plasmodium* infection in the malaria vector *Anopheles*. Here we report the purification and biochemical characterization of CTL4 and CTLMA2 from *An. gambiae* and *An. albimanus*. CTL4 and CTLMA2 are known to form a disulfide-bridged heterodimer via an N-terminal tri-cysteine CXCXC motif. We demonstrate in vitro that CTL4 and CTLMA2 intermolecular disulfide formation is promiscuous within this motif. Furthermore, CTL4 and CTLMA2 form higher oligomeric states at physiological pH. Both lectins bind specific sugars, including glycosaminoglycan motifs with β1-3/β1-4 linkages between glucose, galactose and their respective hexosamines. Small-angle x-ray scattering data supports a compact heterodimer between the CTL domains. Recombinant CTL4/CTLMA2 is functional *in vivo*, reversing the enhancement of phenol oxidase activity in ds*CTL4*-treated mosquitoes. We propose these molecular features underline a common function for CTL4/CTLMA2 in mosquitoes, with species and strain-specific variation in degrees of activity in response to *Plasmodium* infection.

## Introduction

The innate immune response of *An. gambiae* to malaria parasites (genus *Plasmodium*) in an infectious blood meal is a significant factor influencing the prevalence and intensity of infectious mosquitoes in a population^1,2^. Understanding the molecular interactions and mechanisms of *Anopheles* immunity is therefore key to comprehending, predicting, and potentially controlling disease transmission.

*An. gambiae* has a complement-like immune response centered upon thioester-containing protein 1 (TEP1) that effectively targets *Plasmodium* ookinetes following their traversal of the midgut epithelium, prior to their transformation into oocysts^3-7^. The immune response to *Plasmodium* involves additional proteins such as the leucine-rich immune molecule (LRIM) family^8^, CLIP proteases^9^, and other families. Two LRIM family members, LRIM1 and APL1C, form a heterodimeric complex that directly interacts with TEP1 to regulate its anti-*Plasmodium* activity^10-12^. Yet the extent of interactions between these and other immune factors and mechanistic details of the mosquito immune response remain in large part unknown.

The C-type lectin (CTL) fold is the most common binding site for glycans, their special feature being a lectin-bound Ca^2+^ ion in direct coordination with the bound sugar^13-15^. The CTL domain (CTLD), or C-type carbohydrate recognition domain (CRD), consists of ∼130 amino acids with a five-stranded antiparallel β-sheet and two α-helices. Four cysteines within the CTLD form two disulfide bonds that stabilize the fold. The Ca^2+^ binding site lies in a loop between the second and third β-strands. However, many proteins have a CTL fold but lack the Ca^2+^-binding site and do not bind sugars, i.e. they are CTLDs but not C-type CRDs^14^.

CTLs form two groups according to binding preference. CTLs with the sequence EPN in the calcium binding site display a preference for binding mannose-type sugars (equatorial 3’, 4’ OH groups, e.g. fucose, glucose), while CTLs with the sequence QPD prefer galactose-type sugars^13^. Two types of lectins in the immune system are collectins and selectins. Collectins such as mannose-binding protein (MBP) and surfactant protein A and D (SP-A, SP-D) are mannose-type secreted homo-oligomers that bind to pathogen surfaces and trigger innate immune responses such as complement. Selectins are mannose-type cell surface receptors that bind Lewis^A^/Lewis^X^ antigens, via the fucose moiety, that promote adherence of leukocytes to vascular walls in the process of extravasion. Collectins have a second Ca^2+^ binding site while selectins have only one.

The CTL proteins CTL4 and CTLMA2 were first reported to influence the immune response of *An. gambiae* to *P. berghei* infection coincident with that of the first LRIM family member LRIM1^16^. RNAi knockdown of TEP1 (ds*TEP1*)^3^ and LRIM1 (ds*LRIM1*)^16^ results in increased numbers of oocysts in the midgut, indicating a refractory or antagonist phenotype. Knockdown of CTL4 (ds*CTL4*) or CTLMA2 (ds*CTLMA2*) results in significantly reduced numbers of oocysts and increased melanization, implying a susceptible or agonist phenotype.

In the *An. gambiae* L3-5 strain, parasites targeted by TEP1 are killed by lysis, followed by melanization of corpses^17,18^. In the *An. gambiae* G3 strain, melanization requires knockdown of CTL4, in which case it may lead to killing independently of lysis^18^. Melanization in the absence of CTL4 or CTLMA2 required the function of LRIM1. This suggests that CTL4 and CTLMA2 act to suppress either the targeting of *Plasmodium* ookinetes or the downstream melanization response^18^.

CTL4 and CTLMA2 cooperate to protect mosquitoes from infection with Gram-negative bacteria^19^. Either ds*CTL4* or ds*CTLMA2* resulted in decreased survival following infection with *E. coli* but not infection with *S. aureus*. Although ds*CTL4* increased the melanization of *Plasmodium* parasites, it was not observed to increase phenol oxidase activity in the hemolymph following bacterial challenge. Intriguingly, CTL4 and CTLMA2 form a disulfide-bridged heterodimer via an N-terminal CXCXC motif that is necessary for their stability in the hemolymph, analogous to the heterodimer between LRIM1 and APL1C.

The agonist effect of CTL4/CTLMA2 on *P. berghei* was not initially replicated using the human malaria parasite *P. falciparum*^20^. However, this reflects the naturally lower level of infection intensity for the human parasite; increased infection levels resulted in melanization of *P. falciparum* upon CTL4/CTLMA2 knockdown^21^, and most importantly, strain-specific mosquito parasite interactions and the ability of some parasite strains to evade the mosquito immune response^7,22^. The phenotype of CTL4/CTLMA2 silencing also depends on the specific host species. In *An. albimanus* CTL4/CTLMA2 is antagonistic towards both *P. berghei* and *P. falciparum*^21^. This has led to the hypothesis that the functions of these two proteins or associated cofactors, have diverged and their mechanistic involvement in regulating infection intensity and melanization is unlinked.

These data led us to address the question, what is the molecular structure and function of CTL4/CTLMA2 and to what extent is it conserved throughout the *Anopheles* genus? We report that CTL4/CTLMA2 intermolecular disulfide bond formation can occur via any two cysteines of the CXCXC motif, and that the two proteins can form higher-order oligomers via complementary electrostatic interactions. The solution structure of CTL4 and CTLMA2 was determined by small-angle x-ray scattering. Analysis of glycan binding of CTL4 and CTLMA2 suggest the heterodimer separately and synergistically recognize β1-3 and β1-4 glucose/galactose linkages. CTL4 knockdown results in increased PO activity following *E. coli* challenge, which is reversed by co-silencing of TEP1 or co-administration of recombinant CTL4/CTLMA2. Taken together, these results suggest a conserved molecular function of these two lectins in the regulation of melanization downstream of immune recognition in anopheline mosquitoes.

## Results

### Conservation of CTL4/CTLMA2 in Anopheles

CTL4 and CTLMA2 both consist of a signal peptide, a short N-terminal sequence containing the CXCXC motif, and a single CTL domain. A recent report suggested that the function of CTL4 and CTLMA2 or their cofactors have diverged within *Anopheles*, and specifically that *An. albimanus* CTL4 does not contain the N-terminal cysteine residues involved in disulfide linkages^21^. We first reexamined the orthologs of *CTL4* (AGAP005335) and *CTLMA2* (AGAP005334), which have a close back-to-back orientation on chromosome 2L (Fig. 1a), using the current gene models (as of February 2019) in Vectorbase^23^. We found proteins with an N-terminal CXCXC motif for 15/16 CTL4 orthologs, including *An. albimanus* AALB014534, and 13/14 CTLMA2 orthologs. We performed multiple sequence alignments of both CTL4 and CTLMA2 from 10 Asian, African and New World *Anopheles* species (Fig. 1b). The unrooted phylogenetic trees have almost identical topology, and a combined phylogenetic tree has two symmetric branches, except for the position of *An. dirus* with respect to *An. funestus* and *An. maculatus*.

**Figure 1.**
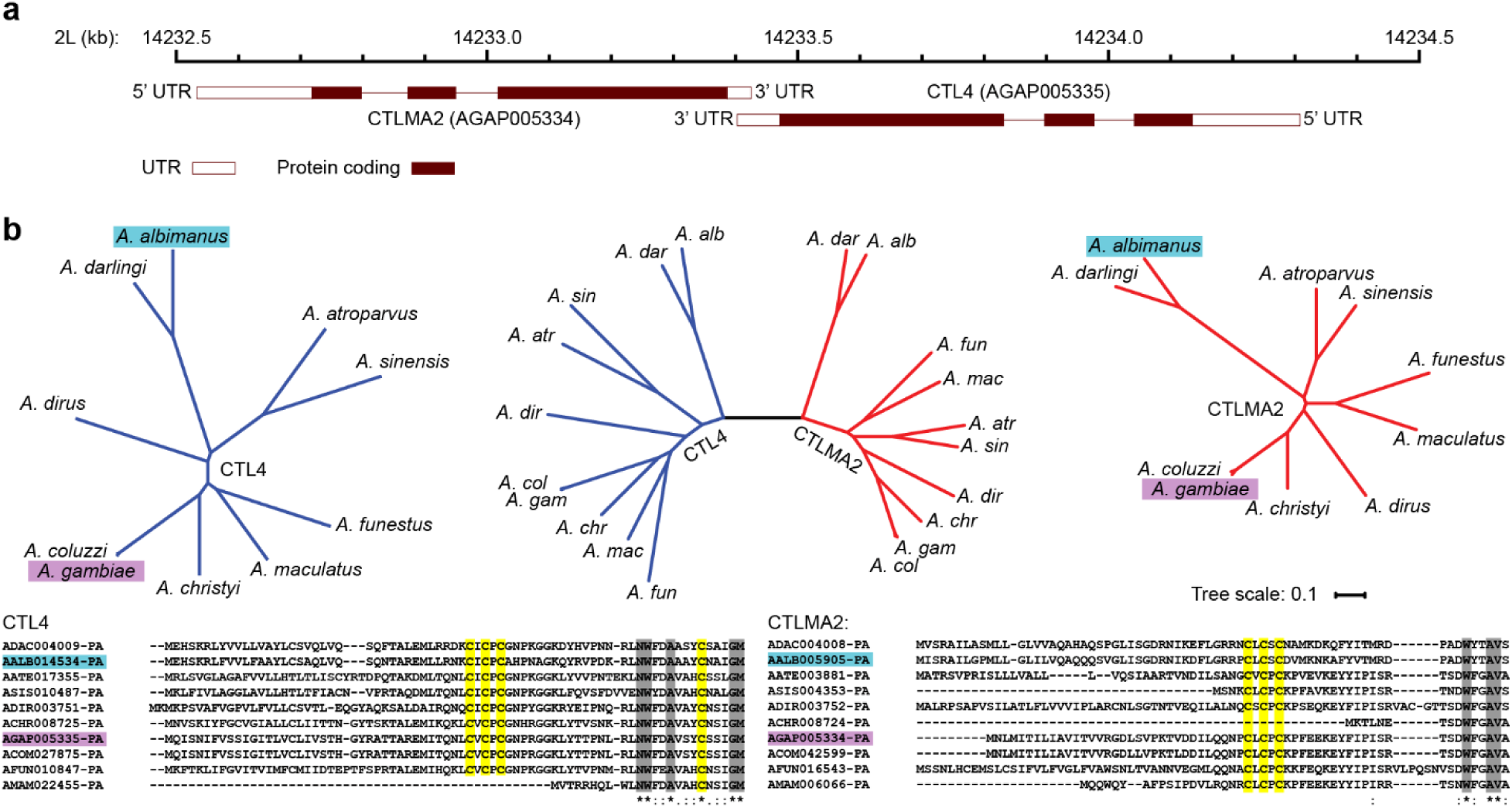
Conservation of *CTL4* and *CTLMA2* in *Anopheles*. (**a**) Schematic illustrating back-to-back arrangement of *An. gambiae* CTL4 and CTLMA2 on chromosome 2L. (**b**) Unrooted phylogenetic trees for CTL4 (blue), CTLMA2 (red) in ten *Anopheles* species including *An. gambiae* (purple) and *An. albimanus* (cyan). The N-terminal portion of both multi-sequence alignments is shown with conserved cysteine residues highlighted yellow, other conserved residues highlighted in gray. The CXCXC motif is conserved in all sequences except those truncated at the N-terminus.

This supports the hypothesis that *CTL4* and *CTLMA2* are conserved within the *Anopheles* genus, with missing orthologs reflecting incomplete annotation of certain genomes. We examined the genomic region of three species with a reported CTL4 ortholog but no *CTLMA2* ortholog: *An. stephensi* ASTE002637, *An. minimus* AMIN007380, and *An. melas* AMEC014491. All possess a close or overlapping gene in inverse orientation – ASTE002636, AMIN007379 and AMEC088499, comprising two CTL domains. For *An. stephensi* and *An. minimus* the second CTL is preceded by a CXCXC motif, suggesting they may be CTLMA2 orthologs. In *Aedes* and *Culex* mosquitoes, the situation is less clear. A *CTLMA2* ortholog including an N-terminal CXCXC motif is annotated in *Aedes albopictus*, AALF001196, and has a close back-to-back inverted two-exon gene AALF001195, comprising of a serine protease and a CTL domain. The October 2018 gene model for *Ae. aegypti* includes a *CTLMA2* ortholog with N-terminal CXCXC motif, CTLMA14 (AAEL014382), currently listed as a CTL4 ortholog). A *CTLMA2* ortholog is annotated in *Culex quinquefasciatus*, CPIJ000443, but lacks the N-terminal CXCXC motif. This suggests that *CTLMA2* arose in a common ancestor of *Anopheles* and *Aedes* whereas *CTL4* may be *Anopheles*-specific.

### Biochemical Characterization

*An. gambiae* CTL4/CTLMA2 (Ag, Fig. 2a,b), CTL4 and CTLMA2 CRDs (Fig. S1a), and *An. albimanus* CTL4/CTLMA2 (Aa, Fig. S1b) were expressed in insect cells using the baculovirus expression vector system (BEVS) and purified to homogeneity. Mature AgCTL4 (25–177) has a molar mass of 17.3 kDa and pI 7.7. Mature AgCTLMA2 (18–174) has a molar mass of 17.9 kDa and pI 4.5. The purified heterodimer has a molecular weight on non-reducing (NR) SDS-PAGE of 35 kDa (Fig. 2a) and elutes as a single peak on size-exclusion chromatography (SEC) with an apparent molecular weight of 40 kDa (Fig. 2b). Monomeric CTL4 and CTLMA2 appear on reducing SDS-PAGE at 17 kDa and 20 kDa, respectively. As previously noted, the increased apparent molecular weight of CTLMA2 may reflect its unusual (acidic) amino acid distribution^19^.

**Figure 2.**
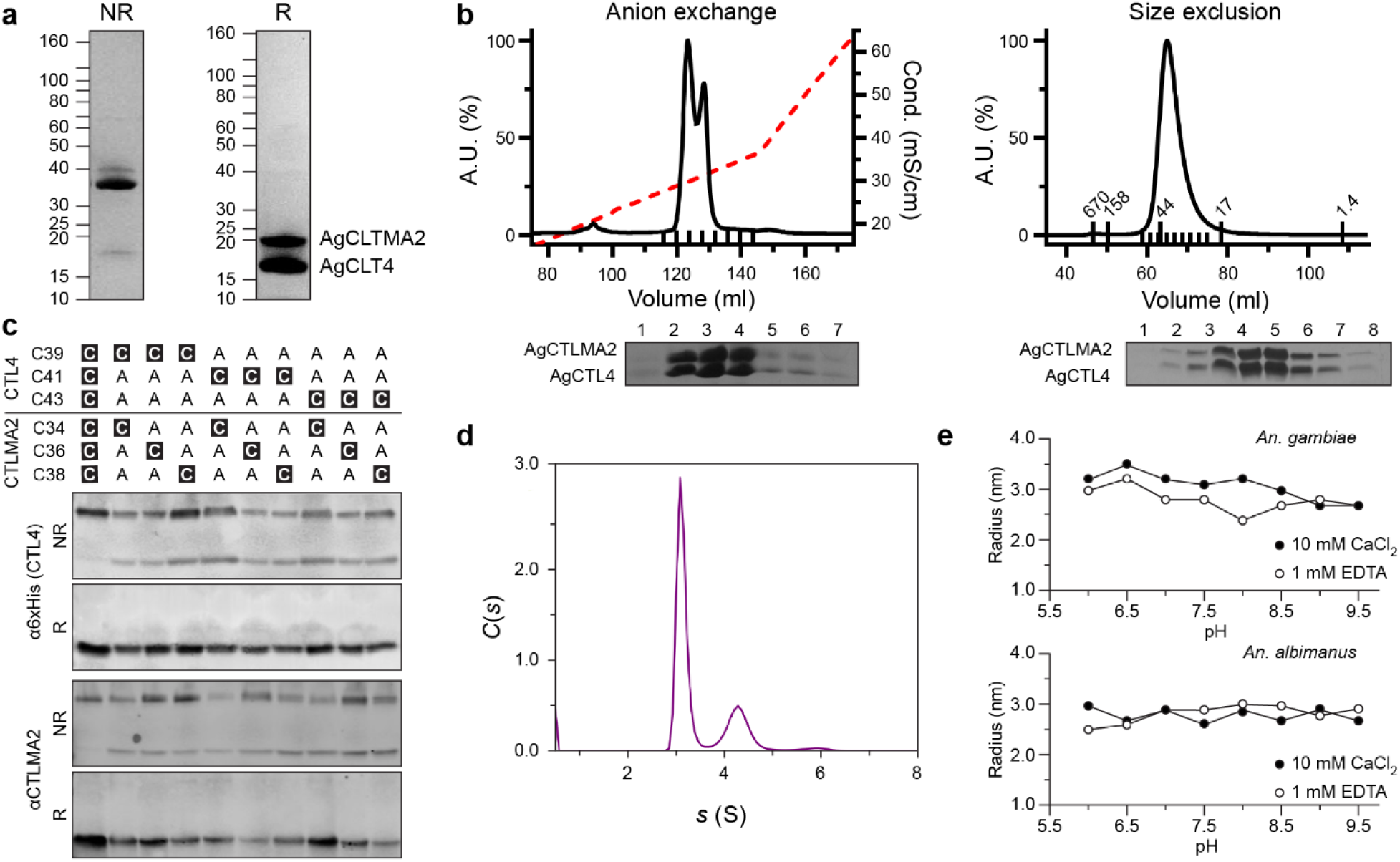
Biochemical characterization of recombinant CTL4/CTLMA2. (**a**) Purified *An. gambiae* CTL4/CTLMA2 heterodimer on non-reducing (NR) and reducing (R) SDS-PAGE. Representative of >10 independent experiments. (**b**) Anion exchange (MonoQ 10/10) and size-exclusion (Superdex75 16/60) chromatogram for *An. gambiae* CTL4/CTLMA2. Small ticks indicate accompanying SDS-PAGE fractions, large ticks indicate SEC MW standards. Representative of >10 independent experiments. (**c**) α6×His (CTL4) and αCTLMA2 Western blotting for nine cysteine mutants of the CTL4/CTLMA2 heterodimer on non-reducing (NR) and reducing (R) SDS-PAGE. In all mutants heterodimer formation is evident, though less efficient. Representative of three independent experiments. (**d**) Plot of *C*(*s*) vs. *s* for sedimentation velocity analytical ultracentrifugation of 1 mg/ml CTL4/CTLMA2, 200 mM NaCl, 20 mM HEPES pH 7.5. Three peaks of increasing sedimentation coefficient are observed, *s*_1_ = 3.1 s, *s*_2_ = 4.4 s, *s*_3_ = 5.9 s. Representative of three independent experiments. (**e**) Dynamic light scattering of CTL4/CTLMA2 vs. pH. Data is fitted to a spherical particle whose radius reflects the size distribution in solution, no trend is apparent in the pH range 6–9.5 for *An. gambiae* or *An. albimanus* CTL4/CTLMA2 in either 1 mM EDTA or 10 mM CaCl_2_. Result from one of two independent experiments.

It has been shown that *An. gambiae* CTL4/CTLMA2 is stabilized by an intermolecular disulfide between cysteines of the N-terminal CXCXC motif; mutation of these three cysteines to alanine abrogates interchain disulfide formation^19^. To further refine the location of the interchain disulfide, we constructed nine mutants containing a single cysteine in the N-terminal CXCXC motif of CTL4 and CTLMA2, co-expressed the proteins in Sf9 cells, and performed Western Blotting to detect CTL4 and CTLMA2. Intermolecular disulfide formation was inefficient but evident on non-reducing SDS-PAGE in all cases (Fig. 2c).

This suggests that N-terminal intermolecular disulfide formation between CTL4 and CTLMA2 is promiscuous, rather than involving a specific cysteine residue from each protein. If intermolecular disulfide formation is promiscuous, CTL4/CTLMA2 heterodimers could in principle form multivalent disulfide-bridged oligomers. No disulfide-linked oligomers are detected on NR-SDS-PAGE for *An. gambiae* CTL4/CTLMA2 (Fig. 2a). Similarly, while CTL4 and CTLMA2 can form disulfide-bridged homodimers *in vitro*^19^, the purified product is overwhelmingly (>95%) heterodimer. Some minor bands (∼10%) are observed on NR-SDS-PAGE for *An. albimanus* CTL4/CTLMA2 (Fig. S1b) that may represent homodimer or oligomer formation.

Both *An. gambiae* CTL4/CTLMA2 and *An. albimanus* CTL4/CTLMA2 are polydisperse in solution. Higher-order non-covalent oligomerization is evident as a shoulder of the main peak in SEC with increasing concentration, and is more pronounced for *An. albimanus* CTL4/CTLMA2 (S1 Fig). We confirmed the existence of oligomeric species by sedimentation-velocity analytical ultracentrifugation (AUC) (Fig. 2d). At pH 7.5 a series of species of decreasing intensity is observed in the *c*(*s*) distribution: *s*_1_ = 3.1 s, *s*_2_ = 4.4 s, *s*_3_ = 5.9 s. Oligomerization is independent of Ca^2+^ for *A. gambiae* CTL4/CTLMA2 but increases substantially with Ca^2+^ for *A. albimanus* CTL4/CTLMA2 (S1B Fig.), and is not correlated with pH according to dynamic light scattering (DLS) (Fig. 2e).

### Calcium binding

As CTLs, both CTL4 and CTLMA2 may bind calcium. However, the canonical Ca^2+^ binding residues in CTL4 are mutated, suggesting it may be a CTLD but not a C-type CRD. Hence, we measured the calcium binding affinity of *An. gambiae* CTL4, CTLMA2 and the CTL4/CTLMA2 heterodimer of *An. gambiae* and *An. albimanus* by isothermal titration calorimetry (ITC). Under equivalent conditions, binding was observed for CTLMA2 and CTL4/CTLMA2, but not for CTL4 (Fig. 3a-c). Binding constants and thermodynamic parameters were calculated from the results of three independent experiments (Table 1). CTLMA2 Ca^2+^ binding is well fit by a single site model with *K*_D_ = 173±27 μM, Δ*H* = 12±2 kcal/mol. The affinity of *An. gambiae* CTL4/CTLMA2 for calcium is ∼40× higher than CTLMA2 with *K*_D_ = 4.9±0.5 μM, Δ*H* = −23±4 kcal/mol. *An. albimanus* CTL4/CTLMA2 (Fig. 3d) has a similar affinity for calcium with *K*_D_ = 2.82 μM, Δ*H* = −12.1 kcal/mol. However, calcium binding to both *An. gambiae* and *An. albimanus* CTL4/CTLMA2 was sub-stoichiometric (*N* = 0.36-0.50).

**Table 1:**
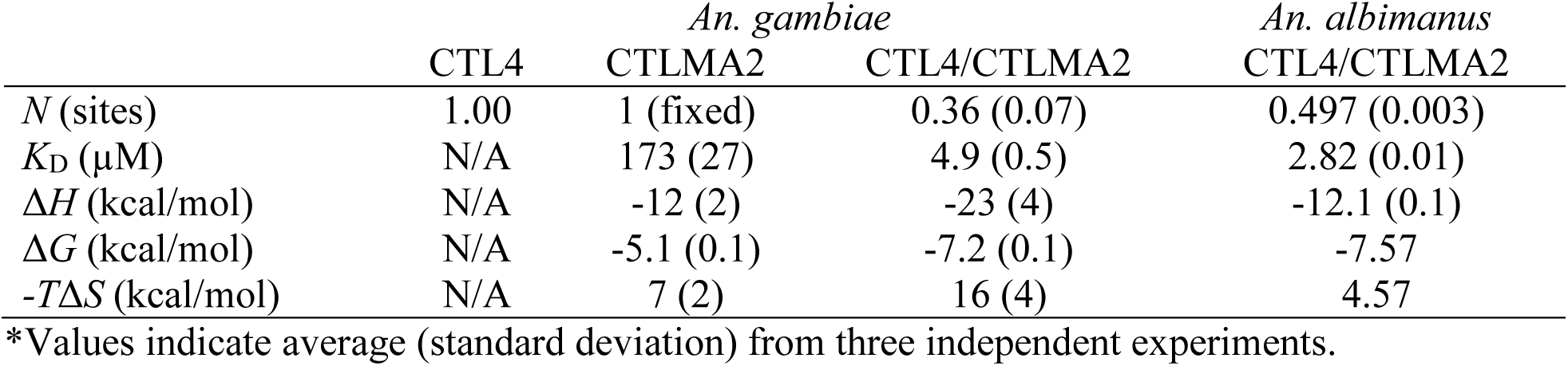
Calcium binding of CTL4 and CTLMA2 by ITC*

**Figure 3.**
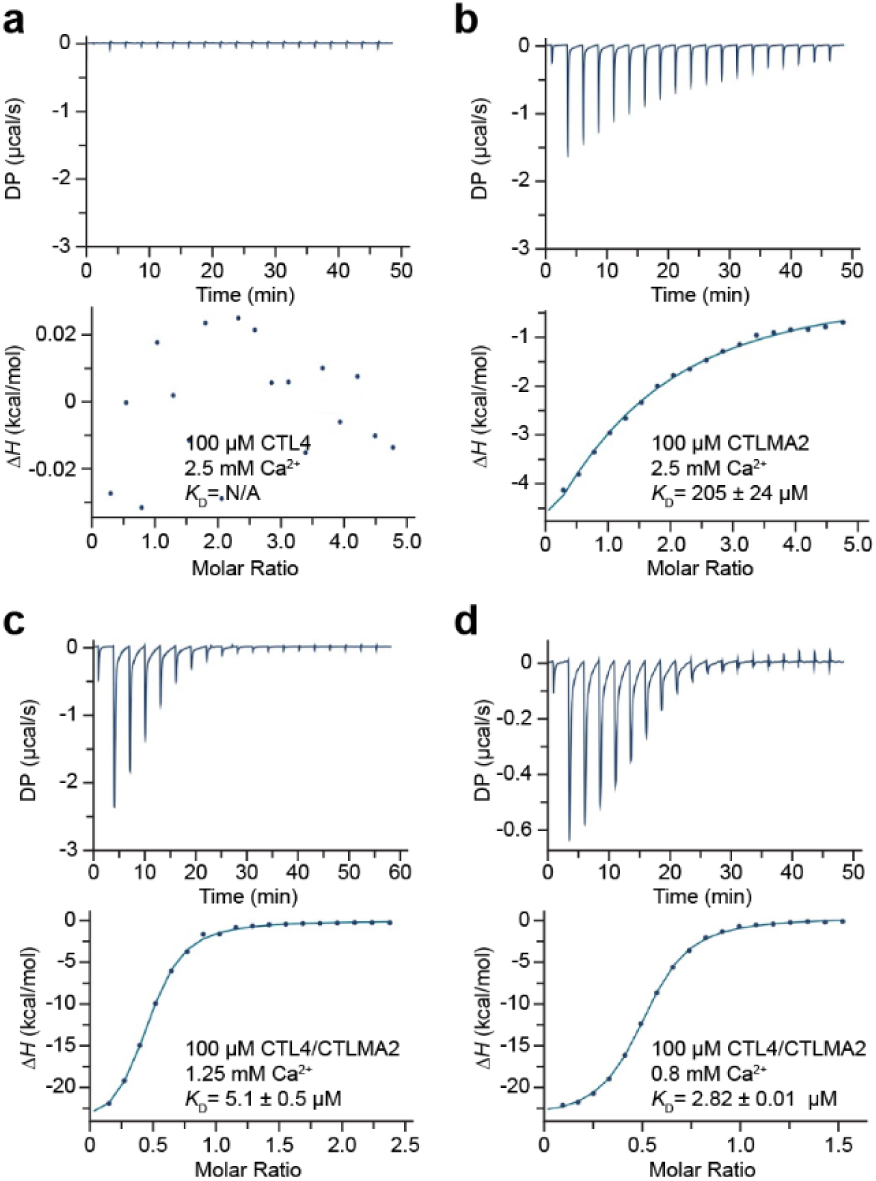
ITC binding isotherm for calcium binding. (**a**) *An. gambiae* CTL4, (**b**) *An. gambiae* CTLMA2, (**c**) *An. gambiae* CTL4/CTLMA2.(**d**) *An. albimanus* CTL4/CTLMA2. Protein concentration in cell was 100 μM, Ca^2+^ (CaCl_2_) concentration in syringe was 2.5 mM for monomers, 1.25 mM *An. gambiae* CTL4/CTLMA2, 0.8 mM for *An. albimanus* CTL4/CTLMA2. No binding is evident for CTL4, weak binding for CTLMA2 and tight binding for CTL4/CTLMA2. Representative of three independent experiments.

### Glycan binding

CTL4 and CTLMA2 belong to the lineage of myeloid CTLs – including macrophage mannose receptor (MMR) and DC-SIGN – that form a conserved family of immune receptors in metazoans^15,24^. Among CTLs with known structure, CTLMA2 has 30% sequence identity to the carbohydrate recognition domain (CRD) of mouse scavenger receptor (SCRL, PDB ID 2OX9)^25^ and porcine surfactant protein D (SP-D, PDB ID 4DN8)^26^. CTLMA2 conserves residues associated with Ca^2+^ binding in the glycan binding loop, and the canonical EPN motif associated with D-mannose selectivity (Fig. 4a). In contrast, CTL4 lacks all residues associated with Ca^2+^ binding, consistent with the fact that no binding was observed by ITC. The only CTL of known structure with considerable similarity to CTL4 is factor IX/X binding protein (X-bp) from the venom of the Chinese moccasin *Deinagkistrodon acutus* (1IOD). X-bp is a modified CTLD in which the glycan binding domain is replaced by a long loop that mediates dimerization to generate the Factor IX/X binding site. Hence, although some insect CTLs do not require calcium for glycan binding, it is unclear if CTL4 should bind glycans at all.

**Figure 4.**
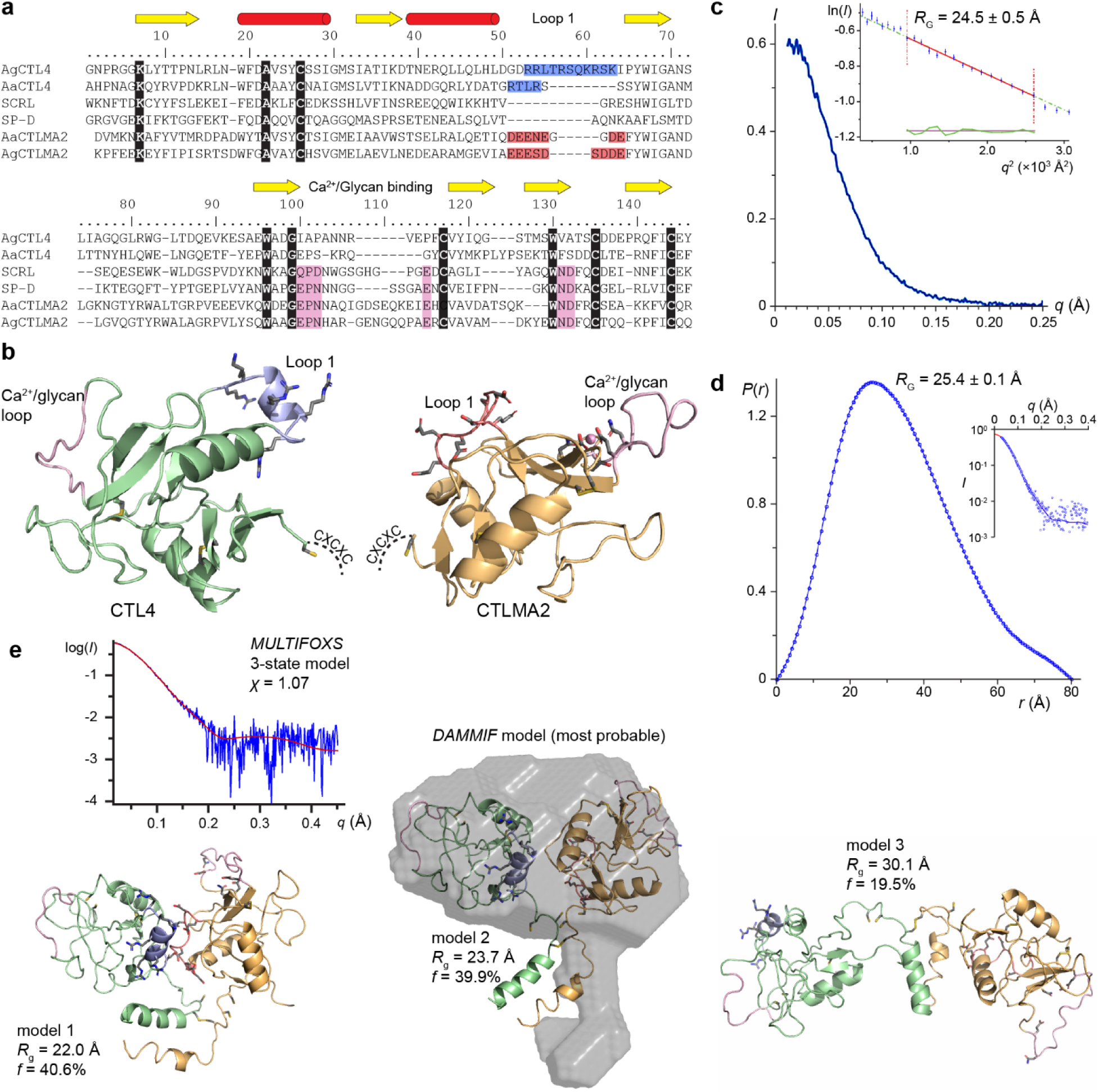
Solution scattering data and model for CTL4/CTLMA2. (**a**) Sequence alignment for *An. gambiae* and *An. albimanus* CTL4 and CTLMA2 with mouse scavenger receptor CTLD (SCRL, PDB ID 2OX9) and porcine surfactant protein D (SP-D, PDB ID 4DN8). Identically conserved residues are highlighted with black. Residues of the Ca^2+^/glycan binding loop are highlighted in pink. CTL4 basic loop 1 residues are highlighted in blue, CTLMA2 acidic loop 1 residues in red. (**b**) Molecular model for CTL4 (green) and CTLMA2 (orange). Ca^2+^/glycan binding loop highlighted in pink. Cysteines in disulfide bonds shown as yellow sticks. Based on proximity of the N-terminal CXC motif to form an intermolecular disulfide, the likely orientation of CRDs in the CTL4/CTLMA2 heterodimer would have outward-facing Ca^2+^/glycan binding loops and inward-facing charged loops. (**c**) Small-angle x-ray scattering curve for *A. gambiae* CTL4/CTLMA2. (inset) Guinier plot (*PRIMUS*), *R*_G_=24.5 Å. (**d**) *P*(*r*) distribution (*GNOM*), *D*_max_ = 80 Å. (inset) fit to the scattering curve, *R*_G_=25.4 Å. (**e**) CTL4/CTLMA2 models arising To confirm the glycan array results and to determine binding preferences for the CTLs, SPR analysis was performed (Table 3). In almost all interactions the glycan array and SPR were in agreement for the presence of interactions with the SPR indicating the glycan array had four false negative results: CTLMA2 monomer with H-antigen, chondroitin sulfate and chondroitin-6-sulfate, and CTL4 monomer with chondroitin-6-sulfate. However, all of the false negative results showed binding on the array with the CTL4/CTLMA2 heterodimer.

In order to define their lectin activity, we analyzed CTL4, CTLMA2 and the CTL4/CTLMA2 heterodimer on glycan arrays that display 367 unique glycan structures^27^. These studies demonstrated binding to a range of glycans (Table 2). The monomers of CTL4 and CTLMA2 demonstrated binding to only four and six glycans, respectively, whereas the heterodimer bound 18 different glycans. There is appreciable difference between ligands bound by the individual monomers and the heterodimer; 3/4 (75%) of glycans recognized by CTL4 and 2/6 (33%) of glycans recognized by CTLMA2 were not recognized by CTL4/CTLMA2.

**Table 2:** Glycan array results for CTL4 and CTLMA2*

| ID# | Glycans | CTL4 | CTLMA2 | CTL4/<br>CTLMA2 |
| --- | --- | --- | --- | --- |
| 6 | GalNAcβ-sp3 | | | 1.14 |
| 47 | 6-H<sub>2</sub>PO<sub>3</sub>Manα-sp3 | | | 1.32 |
| 113 | GlcNAcβ1-3GalNAcα-sp3 | | | 1.16 |
| 163 | 6-O-Su-Galβ1-4GlcNAcβ-sp3 | 1.96 | | |
| 199 | 4,6-O-Su<sub>2</sub>-GalNAcβ1-4-(3-O-Ac)GlcNAcβ-sp3 | | 1.42 | |
| 251 | GlcNAcβ1-4Galβ1-4GlcNAcβ-sp2 | | 1.04 | |
| 287 | 3-O-Su-Galβ1-3(Fucα1-4)GlcNAcβ-sp3 | | 1.12 | 1.14 |
| 368 | Fucα1-2(GalNAcα1-3)Galβ1-4GlcNAcβ-sp3 | | | 1.42 |
| 383 | Galβ1-4GlcNAcβ1-3Galβ1-4Glcβ-sp2 | | | 1.47 |
| 421 | Neu5Acα2-3(GalNAcβ1-4)Galβ1-4Glcβ-sp2 | | | 1.94 |
| 428 | Fucα1-3(Neu5Acα2-3Galβ1-4)6-O-Su-GlcNAcβ-sp3 | | 1.01 | 1.38 |
| 479 | Fucα1-2Galβ1-3GlcNAcβ1-3Galβ1-4Glcβ-sp4 | | | 1.92 |
| 488 | Galβ1-4GlcNAcβ1-3(Galβ1-4GlcNAcβ1-6)GalNAcα-sp3 | | 1 | 1.53 |
| 505 | (GN-M)<sub>2</sub>-3,6-M-GN-GNβ-sp4 | | 1.18 | 1.74 |
| 529 | Neu5Acα2-6(Galβ1-3)GlcNAcβ1-3Galβ1-4Glcβ-sp4 | | 1.09 | 1.49 |
| 533 | (Neu5Acα2-8)<sub>2</sub>Neu5Acα2-3(GalNAcβ1-4)Galβ1-4Glc-sp2 | | | 1.58 |
| 536 | Neu5Acα2-3Galβ1-3GlcNAcβ1-3Galβ1-4Glcβ-sp4 | | | 2.13 |
| 625 | (GlcAβ1-4GlcNAcβ1-3)8-NH<sub>2</sub>-ol | | | 2.14 |
| 18B | Galβ1-3GalNAcβ1-3Galα1-4Galβ1-4Glc | 3.65 | | |
| 13K | (GlcA/IdoAβ1-3(±4/6S)GalNAcβ1-4)<sub>n</sub> (n<250) | | | 2.16 |
| 13M | (GlcA/IdoAβ1-3(±6S)GalNAcβ1-4)<sub>n</sub> (n<250) | | | 2.94 |
| 14I | HA 1600000 da 2.5 mg/ml | 1.48 |  | 1.52 |
| 14K | β1-3Glucan | 2.32 | | |

\*Values indicate fold above background plus three standard errors of the background (average of four technical replicates). Abbreviations: GN – GalNac; HA – haemagglutinin; M – mannose.

**Table 3:** Surface plasmon resonance (SPR) of CTL4/CTLMA2 with glycans

| Glycans (common name) | Glycan array ID | CTL4 (μM) | CTLMA2 (μM) | CTL4/CTLMA2 (μM) |
| --- | --- | --- | --- | --- |
| A blood group trisaccharide | 386 | 49.27±12.2 | 1.77±0.19 | 5.56±2.9 |
| B blood group type 5 | N/A | NCDI* | NCDI | NCDI |
| H disaccharide | 479 | NCDI | 10.39±3.26 | 26.56±6.81 |
| asialo-GM1 | 18B | 15.4±3.99 | NCDI | NCDI |
| LNT | 536 | NCDI | NCDI | 8.22±1.71 |
| LNnT | 383,488 | NCDI | NCDI | 17.62±2.99 |
| GlcNAc monosaccharide | 113,251 | 27.87±11.4 | 29.69±7.91 | 21.91±2.8 |
| Chondroitin sulfate | 13K | NCDI | 0.417±0.1 | 0.404±0.19 |
| Chondroitin-6-sulfate | 13M | 33.54±14.2 | 8.47±2.42 | 5.71±1.2 |
| Sialyl-Lewis A | N/A | NCDI | NCDI | NCDI |
| Sulfo-Lewis A | 287 | NCDI | 0.106±0.06 | 0.095±0.02 |
| Sialyl-Lewis X | 428 | NCDI | 0.619±0.1 | 3.81±1.0 |
| Sulfo-lactosamine | 163 | 0.292±0.15 | NCDI | NCDI |

\*NCDI, no concentration-dependent interaction observed within the range of the instruments detection. Abbreviations: GM1 (monosialotetrahexosylganglioside; LNT – lacto-*N*-tetraose; LNnT – lacto-*N*-neotetraose.

The CTLs did not recognize mannose-containing glycans, with the exception of mannose-6-phosphate by CTL4/CTLMA2, despite the canonical EPN motif present in CTLMA2. Rather, the structures recognized are generally glycosaminoglycan (GAG) motifs comprising β1-3/β1-4 linkages between glucose (Glc), galactose (Gal) and their respective hexosamines GlcNac and GalNac. The Galβ1-4Glc linkage was present in 12/23 (52%) of glycans recognized, including 6/23 (26%) containing Galβ1-4GlcNac and 4/23 (17%) containing the keratan motif Galβ1-4GlcNac β1-3Gal or GlcNac β1-3Galβ1-4Glc.

The array also displayed some preference for polymeric and sulfated glycans. Of the four glycans recognized by CTL4, the strongest hit was for globopentaose (Gb5, Galβ1-3GalNAcβ1-3Galα1-4Galβ1-4Glc); CTL4 also bound HA 160 kDa (GlcAβ1-3Glc*N*Acβ1-4)_n_ and β1-3Glucan (Glcβ1-4Glc)_n_. Three of five glycans recognized by the CTLMA2 monomer were sulfated, and the CTL4/CTLMA2 heterodimer recognized chondroitin sulfate and chondroitin-6 sulfate, hyaluroninc acid (GlcAβ1-4GlcNAcβ1-3)_8_, and HA 160 kDa (GlcAβ1-3Glc*N*Acβ1-4)_n_. These results suggest there are distinct and synergistic effects of heterodimerization on glycan binding.

Both CTLMA2 and the CTL4/CTLMA2 heterodimer recognizes several fucose-containing sugars, including sialyl-Lewis^X^ and sulfo-Lewis^A^ but not sialyl-Lewis^A^. The highest affinity binding was observed to sulfo-Lewis^A^ with binding to the CTLMA2 (106 nM) and CTL4/CTLMA2 (95 nM) complex at a KD of ∼100 nM (Table 3). The highest affinity binding observed for CTL4 was also to a sulfated glycan, sulfo-lactosamine (292 nM). from a 3-state fit to the scattering curve (*MULTIFOXS*). One model is aligned with the most probable of 20 *ab initio* bead models fit to the *P*(*r*) distribution (*DAMMIF*). Measurements derived from a single experiment.

### Structural Analysis

To further probe the structure of CTL4 and CTLMA2, we generated structural models of CTL4 and CTLMA2 (Fig. 4b) using *MODELLER*^28^ with additional manual editing. Both CTL4 and CTLMA2 have an extended loop (loop 1) following the second helix of the CTLD (Fig. 4a) with a high density of complementary charged residues; basic residues for CTL4 and acidic residues for CTLMA2. The complementary electrostatics of these loop 1 residues and their proximity to the N-terminal CXCXC motif suggest this is a potential protein/protein interface within the heterodimer (Fig. 4b), for which a hypothetical model was generated with a single disulfide bond in the CXCXC motif. However, the glycan/Ca^2+^ binding loops are a second potential interface, as observed for the two chains of *D. acutus* X-bp. The alternate hypothesis is that the two CTL domains are independent of one another, with flexible linkers joining them via the intermolecular hypothesis.

To test this hypothesis, we analyzed the solution structure of CTL4/CTLMA2 by small angle x-ray scattering (SAXS). Experiments were conducted in 0.5 M NaCl, 20 mM CHES pH 9.0, 0.5 mM CaCl2, and 1% glycerol to minimize interparticle interactions. In these conditions, the protein displayed a linear relation between extrapolated intensity at zero angle and concentration (*I*_0_ vs. *c*) up to a concentration of 3.1 mg/ml. The buffer-subtracted curve of intensity *I* vs. *q* (Fig. 4c) was submitted to *SAXSMoW2* which yields *R*_G_ = 23.0 Å (*I*_0_=0.61) and molecular weight MW = 39 kDa, only 11% greater than the expected heterodimer MW of 35 kDa^29^. However, the calculated Guinier plot is based on only seven data points; fitting over an extended range (Fig. 4c, inset) yields *R*_G_ = 24.5 Å (*I*_0_ = 0.64), while fitting the pairwise distribution function *P*(*r*) (Fig. 4d) yields an *R*_G_ = 25.4 Å (*I*_0_ = 0.65). The *P*(*r*) distribution fit by *DAMMIF*^30^ with 20 *ab initio* bead models with an average normalized structural discrepancy NSD = 1.0 ± 0.2. The main body of the *ab initio* models are similar in shape to the expected CTL4/CTLMA2 heterodimer.

Additional density extends from the main body of the bead models that can reflect either the N-terminal sequence of both proteins including the CXCXC motif, or a minor population of CTL4/CTLMA2 with a higher radius gyration. To test this hypothesis we performed multi-state modeling of the SAXS profile with the program *MULTIFOXS*^31^. We generated a complete model for CTL4-6xHis/CTLMA2 with an N-terminal coiled-coil terminated by an intermolecular disulfide between CTL4 C39 and CTLMA2 C34. *MULTIFOXS* generated an ensemble of 10,000 variants of the model to compare to the experimental scattering curve. The only flexible residues were CTL4 40–45 and 178–183 (6xHis) and CTLMA2 35-39, with an N-terminal coiled-coil serving as a rigid body connecting the two chains. The best one-state model fit the data with *χ*^2^ = 1.13 and had *R*_G_ = 23.7 Å. The minimum *χ*^2^ = 1.07 was achieved with a 3-state model (Fig. 4e), in which 80% of the scattering is contributed by two compact models with *R*_G_ = 22.0 Å And *R*_G_ = 23.7 Å. This data is consistent with formation of a compact heterodimer between CTL4 and CTLMA2.

### CTL4/CTLMA2 inhibit phenol oxidase activation in response to E. coli

CTL4 and CTLMA2 function as inhibitors of the mosquito melanization response to infection. It was previously reported that CTL4 knockdown did not lead to increased phenol oxidase (PO) activity in response to infection with a mixture of *E. coli* and *S. aureus*^19^. The same study however, found that ds*CTL4* and ds*CTLMA2* mosquitoes were specifically susceptible to Gram-negative bacteria. Hence, we re-examined the effect of CTL4 and CTLMA2 knockdown on hemolymph PO activity in response to only *E. coli* infection (Fig. 5a). At 4 h post-infection with *E. coli*, PO activity was significantly enhanced for ds*CTL4* (*p*=0.02) and ds*CTLMA2* (*p*=0,004) mosquitoes compared to *dsLacZ* controls (Fig. 5b). The average knockdown efficiency for *CTL4* and *CTLMA2* was 93±3% and 89±3%, respectively, with >80% in any single experiment.

**Figure 5.**
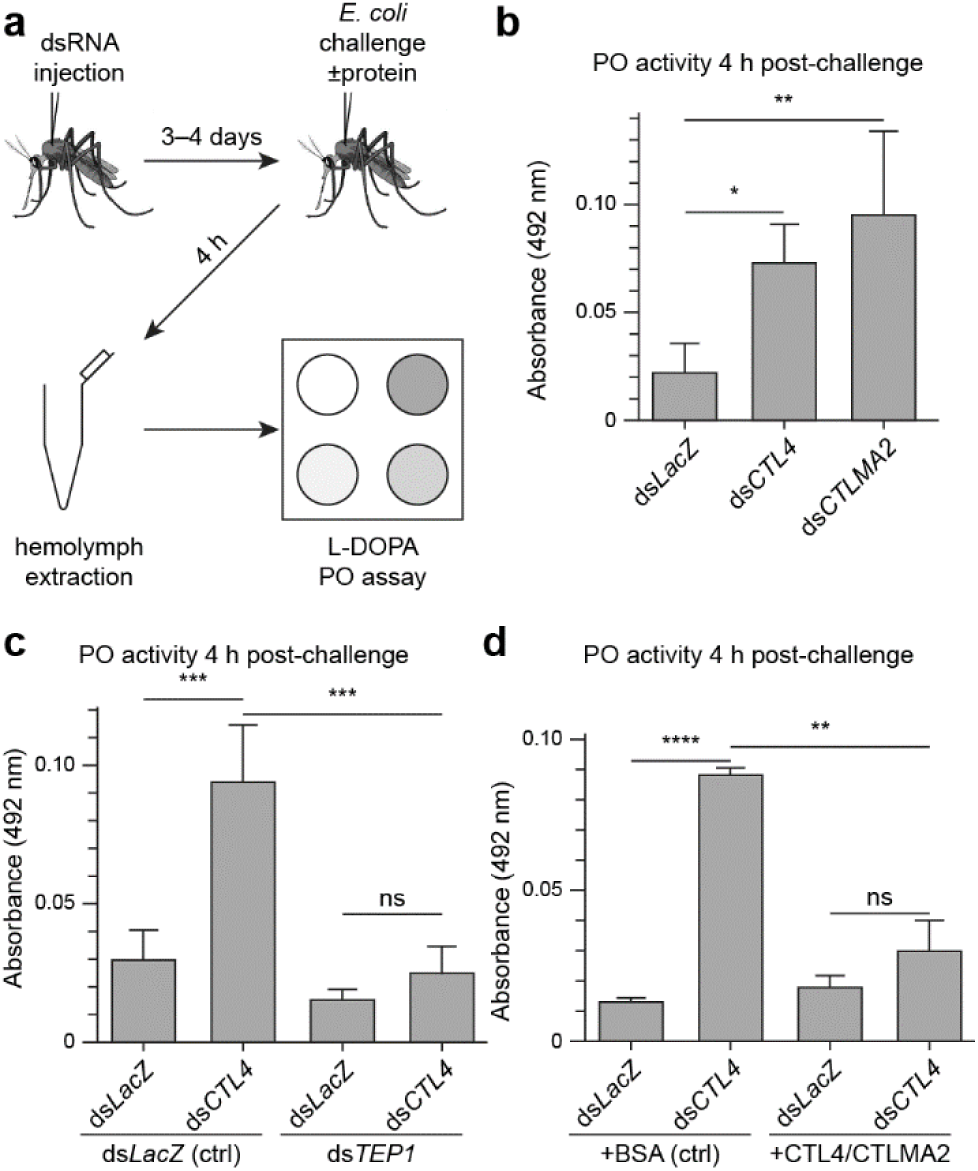
Recombinant CTL4/CTLMA2 inhibits phenoloxidase (PO) activity following *E. coli* challenge (**a**) Experimental design for measuring hemolymph PO activity 4 h after *E. coli* challenge (OD 0.8). PO activity was determined by combining hemolymph with the substrate L-3,4-dihydroxyphenylalanine (L-DOPA) as described in the Materials & Methods. (**b**) Enhanced *E. coli*-induced hemolymph PO activity in ds*CTL4* and ds*CTLMA2* knockdowns. * = 0.017, ** = 0.0035 (*n* = 3, Tukey’s multiple comparisons test) (**c**) Simultaneous knockdown of *TEP1* (ds*TEP1*), compared to *LacZ* control (+BSA), reverses the enhancement of *E. coli*-induced hemolymph PO activity in ds*CTL4* mosquitoes. *** = 0.001 (ds*CTL4*/ds*LacZ* vs. ds*LacZ*/ds*LacZ*), *** = 0.006 (ds*CTL4*/ds*TEP1*vs. ds*CTL4*/ds*LacZ*) (*n* = 3, Tukey’s multiple comparisons test). (**d**) Co-administration of recombinant CTL4/CTLMA2 (+CTLs), compared to BSA control (+BSA), reverses the enhancement of *E. coli*-induced hemolymph PO activity in ds*CTL4* mosquitoes. ** = 0.007, **** = 1.8×10^−5^(*n* = 3, two-tailed heteroscedastic student *t*-test). Bars represent mean ± SD with p ≤ 0.05 considered significant.

Melanization of *Plasmodium* ookinetes upon CTL4/CTLMA2 silencing is dependent on LRIM1^16,21^, TEP1^32^ and SPCLIP1^32^. Since these are all elements of the TEP1 complement-like immune response, we reasoned that enhanced PO activity in the absence of CTL4 should be TEP1-dependent. Accordingly, we compared the enhancement of PO activity in ds*CTL4* mosquitoes with co-administration of ds*TEP1* to co-administration of ds*LacZ* (Fig. 5c). The average knockdown efficiency for TEP1 was 82±9% and >80% in 5/6 experiments (64% in one experiment). Indeed, there was no significant enhancement of PO activity in ds*CTL4* mosquitoes when *TEP1* was also silenced. This confirms that melanization in the absence of CTL4/CTLMA2 is TEP1-dependent.

Co-administration of recombinant CTL4/CTLMA2 with *E. coli* significantly reversed the enhancement of PO activity in ds*CTL4* mosquitoes compared to BSA (*p*=0.007, Fig. 5d). These results demonstrate that CTL4/CTLMA2 is directly involved as a negative regulator of PO activity. In ds*LacZ* mosquitoes however, CTL4/CTLMA2 did not suppress *E. coli*-induced PO activity compared to bovine serum albumin (BSA). Also, *E. coli*-induced PO activity in ds*CTL4* mosquitoes co-injected with recombinant CTL4/CTLMA2 (ds*CTL4*+CTLs) was higher than that of ds*LacZ* mosquitoes co-injected with BSA (ds*LacZ*+BSA). Hence, the rescue effect of injecting recombinant CTL4/CTLMA2 is incomplete compared to the presence of endogenous protein.

## Discussion

Here we describe the biochemical characterization of two C-type lectins, CTL4 and CTLMA2, that play key roles as negative regulators of the melanization cascade in the malaria vector *An. gambiae*. This is relevant to malaria transmission as knockdown of CTL4 and CTLMA2 results in reduced susceptibility to *P. berghei* and, with comparable infection levels, *P. falciparum*. The effect of CTL4 and CTLMA2 on *Plasmodium* is species-dependent, suggesting divergent evolution of the trait during speciation within the *Anopheles* genus. Here we focus on the common molecular features of these proteins conserved throughout *Anopheles*.

Heterodimerization of CTL4 and CTLMA2 is mediated by promiscuous intermolecular disulfide bonding of an N-terminal CXCXC motif conserved throughout *Anopheles* and, for CTLMA2, the *Aedes* genus. Aside from Zn-binding sites in DNA/RNA binding proteins, the CXCXC motif is uncommon in proteins of known structure. At time of publication the motif is present in only 74 representative protein-only structures at <4.0 Å resolution in the Protein Databank. The sequences are generally linear with disulfide bonds (if any) to different strands. Angepoietin-1 has an internal disulfide bond between the first and second cysteines of a CXCXC motif, with the backbone carbonyls serving as ligands to a calcium binding site. This is not observed for CTL4/CTLMA2 since we only observe a single Ca^2+^ binding site by ITC corresponding to the CTLMA2 Ca^2+^/glycan loop.

Steric hindrance tends to limit the formation of multiple disulfide between adjacent strands of closely spaced cysteine residues. However, intramolecular disulfides involving a CXC motif are rare but known within proteins. When CTL4 and CTLMA2 were restricted to a single cysteine within the CXCXC motif, neither protein had a reproducible preference for a single cysteine comparing multiple independent experiments. However, it may still be that a specific intermolecular disulfide is preferred in the context of the wild-type heterodimer produced *in vivo*.

Oligomerization of C-type CRDs is a common feature of CTLs in the immune system^13,14^. The collectins MBP, SP-A, and SP-D form trimeric structures mediated by an N-terminal cysteine-rich region followed by a collagen-like domain. DC-SIGN tetramerizes on cell surfaces via an extended neck region^33-35^. We see no evidence of higher order disulfide-mediated oligomerization of the CTL4/CTLMA2 heterodimer. However, there is clear evidence that CTL4 and CTLMA2 form non-covalent higher-order oligomers by SEC and AUC, especially so for *An. albimanus* CTL4/CTLMA2 in high calcium concentrations (10 mM). The constant ratio of sedimentation coefficients measured by AUC *s*_2_/*s*_1_ = *s*_3_/*s*_2_ = 1.4 is consistent with serial (1, 2, 4, …) oligomerization but no details of the structure or mechanism of this association is yet determined.

The Ca^2+^/glycan binding loop is mutated in CTL4, which does not bind Ca^2+^ but may play a role in oligomerization as observed for the dimerization loop of X-bp. For CTLMA2 Ca^2+^ affinity is enhanced in the presence of CTL4 suggesting a physical interaction either directly or allosterically alters its structure. The CTL4/CTLMA2 Ca^2+^ binding curve is best fit by substoichiometric binding. This could results from residual calcium either incompletely removed by EDTA or reintroduced before the ITC measurement. Or, the Ca^2+^ binding site in CTLMA2 could be partially occluded by misfolding, presence of another divalent metal, oligomerization, interaction with CTL4 or perhaps bound glycans carried over during purification. The observed polydispersity of the recombinant heterodimer may contribute to these experimental factors.

CTL4 and CTLMA2 display several properties analogous to those of collectins. They are oligomeric serum proteins with a protective phenotype vs. Gram negative bacteria. Yet CTLMA2 displays the binding attributes of a selectin. The second calcium binding site in MBP, SP-D, and DC-SIGN is disrupted by mutation of conserved Asn/Asp residues (MBP D188, D194) to His and Arg (CTLMA2 H132, R144). CTLMA2 also binds Lewis^A^/Lewis^X^ antigens as do selectins; the similar affinity of CTL4/CTLMA2 for Lewis^A^/Lewis^X^ suggests this interaction solely involves CTLMA2. CTLMA2 binding of sialylated and sulfated Lewis structures likely involves direct ligation of fucose to calcium, as illustrated by the selectin-like mutant of MBP^36^ and DC-SIGN^37,38^.

Our prior hypothesis was that CTL4, having a mutated Ca^2+^/glycan binding loop, would not recognize glycans. Yet CTL4 recognized a number of specific glycans, including submicromolar affinity for sulfo-lactosamine. Since CTL4 has no Ca^2+^ binding site the mechanism for binding is non-canonical; and notably binding is not conserved in the CTL4/CTLMA2 heterodimer. CTL4 bound GlcNac monosaccharide and chrondoitin-6 sulfate ((GlcA/IdoAβ1-3(±6S) GalNAcβ1-4) with similar affinity to CTLMA2 and the CTL4/CTLMA2 heterodimer. Yet chondroitin sulfate (GlcA/IdoAβ1-3(±4/6S)GalNAcβ1-4) is bound with higher affinity and appears to be mediated solely by CTLMA2. Most intriguingly, binding to lacto-*N*-tetraose (LNT) and lacto-*N*-neotetraose (LNnT) was specific to the CTL4/CTLMA2 heterodimer, suggesting a cooperative binding mechanism.

Binding of polymeric GAGs by CTL4 and CTLMA2 may be relevant to their physiological role as inhibitors of melanization. Insect connective tissues are rich in acidic and neutral GAGs (a.k.a. mucopolysaccharides, mucins), which serve to cement hemocytes together to encapsulate parasites and foreign bodies in the hemocoel^39^. CTL4/CTLMA2 may line connective tissues in mosquitoes, inhibiting self-melanization in response to injury or infection. Such a function is analogous to association of vertebrate Factor H with sialylated surfaces, which serves to inhibit complement activation. Chondroitin and heparin sulfate are present in anopheline mosquitoes and are utilized by *Plasmodium* parasites during invasion^40,41^. If so, we speculate that pathogens, perhaps *Plasmodium*, could recruit glycosaminoglycans or CTL4/CTLMA2 directly to their surface as a means of inhibiting melanization by the host.

A number of questions remain unanswered. Does CTL4/CTLMA2 interact with specific proteins as well as glycans, and if so are such interactions glycan-dependent? Does CTL4/CTLMA2 inhibit melanization even after PO is activated, or does it simply inhibit the proteolytic activation of PO or an upstream factor? Is the binding of protein cofactors dependent on their conformation or proteolytic activation? The ability to reverse enhanced PO activity in ds*CTL4* mosquitoes with co-administration of CTL4/CTLMA2 allows these questions to be addressed by site-specific alterations to the recombinant protein. Elucidating the mechanism of CTL4/CTLMA2 in the melanization response to infection can ultimately address its role in the natural susceptibility or refractoriness of different *Anopheles* species to *Plasmodium* infection.

## Methods

### Protein Expression and Purification

Full-length *An. gambiae* CTL4 (AGAP005335) and CTLMA2 (AGAP005334), *An. albimanus* CTL4 (AALB014534) and CTLMA2 (AALB005905) were obtained by total gene synthesis and subcloned into pFastbac1 with C-terminal 6×His tag For heterodimeric CTL4/CTLMA2 the genes were cloned into pFastbac-Dual with a C-terminal 6×His tag on CTL4. Additional constructs were cloned into pFB-GP67-Hta with a TEV-cleavable 6×His tag. Insect cell lines (sf9 and *T.ni*, Expression Systems LLC) were cultured in ESF-921 (Expression Systems LLC) at 27 °C. Recombinant baculovirus was produced in sf9 cells.

Proteins were expressed in *T.ni* cells, with conditioned media (CM) harvested at 48 h post-infection (hpi). CM was concentrated and diafiltrated into 0.25 M NaCl, 50 mM Tris-HCl pH 7.8 by tangential flow filtration (Centramate™, Pall Biosciences), followed by affinity chromatography using Co Talon resin (Clontech). Further purification was accomplished by ion exchange and size-exclusion chromatography on an AKTA PURE system (GE Healthcare).

### Western Blotting

Sf9 cells were infected at MOI of 0.1 and conditioned media collected 96 hpi. For non-reducing SDS-PAGE DTT was excluded from the 6× loading buffer; samples were not heated prior to electrophoresis on 4–20% mini protean TGX precast gels (BioRad). Gels were transferred to Odyssey nitrocellulose membrane (LI-COR) at 30 V overnight in Towbin buffer with 10% methanol; but 1% SDS in the electrophoresis buffer was necessary for efficient transfer of the heterodimer in non-reducing conditions. Western blotting was performed with anti-6×His mouse mAb (Clontech), with secondary IRDye 800CW goat anti-mouse (LI-COR) for. *An. gambiae* CTL4.

For detection of CTLMA2, full length His-tagged CTLMA2 was expressed in sf9 cells and purified from conditioned media by Co (Talon) affinity chromatography. The protein was used for inoculation into a rat for production of polyclonal antibodys by Cocalico Biologicals (Stevens, PA). The resulting antiserum was affinity purified using recombinant CTLMA2 immobilized on Aminolink resin (Thermo Fisher). Western blotting was performed with affinity purified anti-CTLMA2, with secondary IRDye 680LT goat anti-rat (LI-COR).

### Analytical ultracentrifugation (AUC)

Sedimentation velocity AUC was performed on a Beckman XL-I ultracentrifuge with absorbance optics set at 280 nm. First *An. gambiae* CTL4/CTLMA2 was analyzed under same conditions as for SEC (Fig. 2d). To test effect of calcium (Fig. S1b), *An. gambiae* and *An. albimanus* CTL4/CTLMA2 was prepared at 0.5 mg/ml in 0.15 M NaCl, 20 mM Tris pH 7.5 and either 10 mM CaCl_2_ (Ca-TBS) or 1 mM EDTA (EDTA-TBS). Protein partial specific volume and buffer viscosity were calculated with *SEDNTERP*: 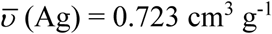, 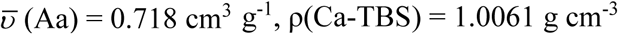, ρ(EDTA-TBS) = 1.0054 g cm^−3^, η(TBS) = 1.002 cP. Samples were centrifuged at 20 °C, 40,000 rpm. Data analysis was performed with *SEDFIT*^42^.

### Dynamic Light Scattering (DLS)

DLS experiments were performed on a DynaPro Plate Reader (Wyatt Technologies) using the Solution and Stability Screen 2 (Hampton Research). CTL4/CTLMA2 in 0.2 M NaCl, 20 mM HEPES pH 7.5 was diluted to 1 mg/ml in a final volume of 20 μl and 4× dilution of the reagents A2–12, H2–H12 resulting in 50 mM buffer ± 1.0 M NaCl.

### Isothermal Titration Calorimetry

Binding studies were performed at 25 °C with a MicroCal PEAQ-ITC (Malvern Analytical). Purified proteins were dialyzed against TBS buffer (0.15 M Nacl, 20 mM Tris, pH 8.0) containing 5 mM EDTA to remove bound calcium. Proteins were subsequently dialyzed in Chelex (Biorad) treated TBS buffer (pH 7.5) to remove both EDTA and calcium. Final protein and ligand solutions were made in the same Chelex-treated TBS (pH 7.5) and filtered through 0.2 μm before loading. The cell was loaded with 300 μl of 100 μM protein. The syringe contained 2.5 mM CaCl_2_ for CTL4 and CTLMA2, 1.25 mM CaCl_2_ for CTL4/CTLMA2. Titrations comprised 19 injections – 2 μl first injection followed by 18× 4 μl injections – with 150 s between each injection. A TBS buffer control was titrated against ligand and used as reference. Integrated heats were fit with a single-site binding model – fixed *N*=1 for CTLMA2, floating *N* for CTL4/CTLMA2 – to determine dissociation constant *K*_D_ and thermodynamic parameters. The results for *An. gambiae* samples arise from three independent experiments. For *An. albimanus* CTL4/CTLMA2 three experiments were performed but under different conditions, with a range of resulting *K*_D_ from 0–5 μM and Δ*H* from 10–27 kcal/mol; The result reported corresponds to equivalent conditions as for *An. gambiae* CTL4/CTLMA2 and with median calculated values for *K*_D_, Δ*H*.

### Analysis of glycan binding by glycan array

Glycan arrays were produced and the glycan library used for screening as previously described^27^. Results were derived from a single batch of each protein, values are the average of four technical replicates. Glycan arrays were probed with 1 µg of 6×His-tagged CTL4, CTLMA2 and CTL4/CTLMA2 in PBS containing 1 mM CaCl_2_ and 1 mM MgCl_2_. A mouse anti-His primary antibody was added (1:1 molar ratio with protein) followed by rabbit anti-mouse Alexa647 (1:0.5 ratio with primary), and goat anti-rabbit Alexa647 (1:0.5 ratio with secondary). The final volume applied to the array was 300 µl incubated for 15 min. Arrays were washed 3 times in PBS containing 1 mM CaCl_2_ and 1 mM MgCl_2_. Arrays were scanned on an Innoscan 1100AL using the 632 nm laser. Binding was considered to be positive when values were 3 standard deviations above the background of the array.

### Analysis of glycan binding by surface plasmon resonance (SPR)

SPR was performed on a Biacore T200 using a CM5 senor chip. Flow cell one was left blank with ethanolamine only blocking the NHS activated carboxydextran. The proteins were immobilized onto flow cell 2-4 (CTL4, CTLMA2, CTL4/MA2 respectively). Glycans were flowed from 160 nM to 100 µM across a 1:5 dilution. A 15 s enhancement injection of PBS containing 1 mM CaCl_2_ and 1 mM MgCl_2_ preceded the sample injection and a 60 second regeneration injection of 10mM Tris 1mM EDTA followed the sample injection. Analysis was performed using the Biacore T200 evaluation software using the surface bound menu; Affinity; Steady state affinity. Glycan were run in triplicate with glycans run in single replicates with the program repeated three times.

### Small-Angle X-ray Scattering

SAXS data was collected in-house on a BioSAXS-2000 (Rigaku Corp.) with a DECTRIS PILATUS 100K detector. Samples were prepared as a dilution series from 1–4.5 mg/ml in 0.5 M NaCl, 20 mM HEPES pH 9.0, 0.5 mM CaCl_2_, 1% glycerol. Data acquisition was 30 min collected as 5 min exposures to ensure no measurable radiation-induced changes within the aquisition period. Following conversion to *I* vs. *q* curves, primary data analysis using the *ATSAS* software package^19,43^. Buffer subtraction and Guinier analysis was performed with *PRIMUS*, *P*(*r*) calculation with *GNOM*. Construction of *ab initio* bead models were derived for each structure was performed with *DAMMIF* using *D*_max_=80 Å, P1 symmetry.

### Mosquito rearing and maintenance

The *An. gambiae* G3 strain was obtained through BEI Resources, NIAID, NIH: *An gambiae*, Strain G3, MRA-112, contributed by Mark Q. Benedict. Mosquitoes were reared on a 12 hr light/dark cycle at 28°C and 75% relative humidity. Larvae were provided fish flakes (Tetramin) and dry cat food (Friskies) until pupation. Adults were maintained on 10% sucrose and fed sheep blood (HemoStat Laboratories, #SBH100) for egg production. Experiments were performed with 2–3 d old adult femailes from independent generations.

### Gene silencing by RNAi

T7 promoter-tagged templates for dsRNA synthesis were generated from a clone in plasmid pIB (*LacZ*), a clone in plasmid pIEx-10 (*CTLMA2*), and from mosquito cDNA (*CTL4* and *TEP1*) using the iProof High-Fidelity PCR Kit (Bio-Rad, #1725331) and purified with the GeneJET PCR Purification Kit (Thermo Fisher Scientific, #K0701) according to the manufacturer provided instructions. Double stranded RNA reactions were performed using the HiScribe T7 High Yield RNA Synthesis Kit (NEB, #E2040S) and purified with the GeneJET RNA Purification Kit (Thermo Fisher Scientific, #K0731) according to the manufacturer’s instructions. Double stranded RNA was reconstituted in ultrapure water to 3 μg/ul for microinjection. For single and double gene knockdown experiments, mosquitoes were injected with 69 or 138 nl dsRNA, respectively. Experiments were performed 3–4 d after dsRNA injection. Gene silencing efficiencies were performed as described previously^18^ with the following modifications: cDNA was synthesized from 1 μg total RNA using the iScript cDNA Synthesis Kit (Bio-Rad, #1708891) following the manufacturer’s instructions and qRT-PCR was performed with a QuantStudio 6 Flex Real-Time PCR System (Applied Biosystems) using PerfeCTa SYBR Green SuperMix, Low ROX (Quanta BioSciences, Inc. #95056-500). T7 template and qPCR primer pair sequences are provided as Supporting Information (Table S2).

### Phenol oxidase (PO) activity assay

PO activity was measured in mosquito cohorts 4 h after the injection of *E. coli* strain DH10B or a mixture of *E. coli* and protein. Mid-log phase *E. coli* was rinsed and resuspended in PBS to OD 0.8 (Average dose: 124,063 CFU/μl). For PO assays involving the co-administration of *E. coli* and protein, bovine serum albumin (BSA, Sigma #B2518) or recombinant *An. gambiae* CTL4/CTLMA2 was combined with *E. coli* to a final concentration of 2.5 µg/µl just prior to injection. Mosquito injections, hemolymph collection, protein quantification, and the PO activity assay were performed as described previously^19^ with the following modifications: (1) For PO assays following the injection of *E. coli* or a mixture of *E. coli* and protein, PO activity was assessed using 4-5 µg of hemolymph protein or the total hemolymph protein obtained from 100 mosquitoes, respectively. (2) Absorbance at 492 nm was recorded every 10 mins for 1 hour in a Molecular Devices SpectraMax 190 plate reader.

## Acknowledgements

This work was supported in part by the National Institute of General Medical Sciences R01GM114358 of the National Institutes of Health to RHGB, National Institute for Allergy and Infectious Diseases R01AI139060 of the National Institutes of Health to MP, and National Health and Medical Research Council (NHMRC; Australia) Program Grant 1071659 and Principal Research Fellowship 1138466 to MPJ.

## Contributions

R.B. and G.L.S. contributed equally to this work. Conceptualization, R.B., A.C., G.L.S., M.P.J., M.P. and R.H.G.B.; Methodology, R.B., A.C., C.J.D., D.C.F.H., L.A.P., G.L.S., M.P.J., M.P., A.M.V. and R.H.G.B.; Investigation, R.B., A.C., C.J.D., D.C.F.H., L.A.P., D.S. and G.L.S.; Visualization, R.B., A.C., C.J.D., L.A.P., G.L.S. and R.H.G.B.; Writing – Original Draft, R.H.G.B.; Writing – Review and Editing, C.J.D., G.L.S., M.P.J., A.M.V., M.P. and R.H.G.B.; Funding Acquisition, M.P.J., M.P., A.M.V. and R.H.G.B.; Supervision, M.P.J., M.P., A.M.V. and R.H.G.B.

## Data Availability

The small-angle x-ray scattering dataset generated analysed in the current study is available in the SASBDB repository as entry SASDFL4, https://www.sasbdb.org/data/SASDFL4.

