## Supplementary Information for "Solution structure, glycan specificity and of phenol oxidase inhibitory activity of *Anopheles* C-type lectins CTL4 and CTLMA2"

**Table S1: dsRNA and qPCR primers used**

| Name | Sequence |
|---|---|
| *T7 Primers* | |
| LacZ-T7 F | taatacgactcactatagggAGAATCCGACGGGTTGTTACT |
| LacZ-T7 R | taatacgactcactatagggCACCACGCTCATCGATAATTT |
| AgCTL4-T7 F | taatacgactcactatagggTGGTTTGATGCCGTGTCCT |
| AgCTL4-T7 R | taatacgactcactatagggAATAAATTGTCTCGGTTCATCATC |
| AgCTLMA2-T7 F | taatacgactcactatagggGCCTTTGCCCGTGCAAACCGTTC |
| AgCTLMA2-T7 R | taatacgactcactatagggTTGACAGATGAACGGTTTCTGCTG |
| AgTEP1-T7 F | taatacgactcactatagggTTTGTGGGCCTTAAAGCGCTG |
| AgTEP1-T7 R | taatacgactcactatagggACCACGTAACCGCTCGGTAAG |
| *qPCR primers* | |
| AgS7-qPCR F | GTGCGCGAGTTGGAGAAGA |
| AgS7-qPCR R | ATCGGTTTGGGCAGAATGC |
| AgCTL4-qPCR F | GCACGGGTACAGGGCTACTA |
| AgCTL4-qPCR R | GCGTGGTGTACAGCTTTCCT |
| AgCTLMA2-qPCR F | GCTGTCATCACAGTGGTTCG |
| AgCTLMA2-qPCR R | GGGTTTTGTTGAAGAATATCATCC |
| AgTEP1-qPCR F | AAAGCTGTTGCGTCAGGG |
| AgTEP1-qPCR R | TTCTCCCACACACCAAACGAA |

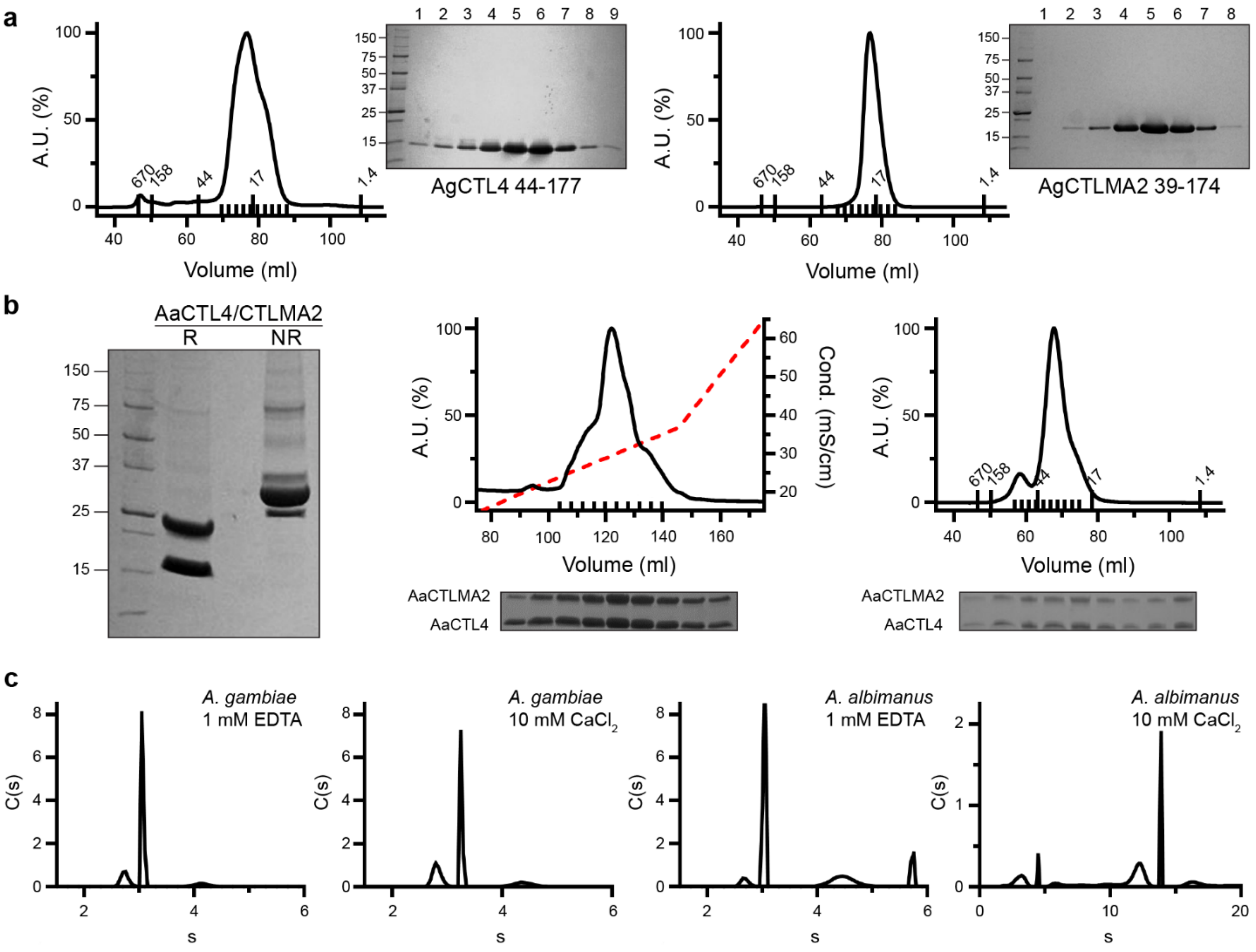


**Figure S1**. Purification of *An. gambiae* CTL4, CTLMA2 and *An. albimanus* CTL4/CTLMA2. (**b**) Reducing (R) and non-reducing (NR) SDS-PAGE, anion exchange (MonoQ 10/10) and size-exclusion (Superdex75 16/60) chromatogram for *An. albimanus* CTL4/CTLMA2. Small ticks indicate reducing SDS-PAGE peak fractions, large ticks indicate SEC MW standards. Representative of >5 independent experiments. (**b**) Plot of *C*(*s*) vs. *s* for sedimentation velocity analytical ultracentrifugation of 0.5 mg/ml *An gambiae* and *An. albimanus* CTL4/CTLMA2, in 0.15 M NaCl, 20 mM Tris pH 7.5 and either 10 mM $CaCl_2$ (Ca-TBS) or 1 mM EDTA (EDTA-TBS). Results from a single experiment.
